# PRPF8-associated retinitis pigmentosa variant induces human neural retina-autonomous photoreceptor defects

**DOI:** 10.1101/2025.06.04.657822

**Authors:** Felix Zimmann, Poulami Banik, Jan Kubovčiak, Prasoon K. Thakur, Martin Čapek, Michal Kolář, Eva Hrubá, Robert Dobrovolný, Zuzana Cvačková, Tomáš Bárta, David Staněk

## Abstract

Retinitis pigmentosa (RP) is an inherited retinal disorder characterized by the progressive loss of photoreceptors that currently lacks effective treatment. Here, we investigated the impact of the pathogenic PRPF8-Y2334N variant on neural retina cells in hiPSC-derived retinal organoids. Expression of this variant resulted in photoreceptor defects, including thinning of the outer segment layer. At the molecular level, we observed relatively minor changes in mRNA expression in multiple retinal cells, indicating that neural retina cells are impacted independently of retinal pigment epithelium (RPE). We found splicing alterations in genes associated with neural and retinal diseases, including those involved in intraflagellar transport, suggesting that these genes may represent common targets of splicing factor mutations. We further detected the misexpression of several circular RNAs (circRNAs), which could serve as early biomarkers of splicing defects caused by RP mutations. Together, we present a model of RP that recapitulates photoreceptor degeneration and demonstrates that these defects are independent of RPE erosion.

## Introduction

Retinitis pigmentosa (RP) is the leading cause of inherited blindness, with a prevalence of 1:4000 (1–3). This disorder is characterized by a progressive degeneration of the retina with the loss of the rod photoreceptor cells preceding the loss of cone photoreceptor cells (2). The primary retinal cells involved are the photoreceptor cells and the cells of the retinal pigment epithelium (RPE) (4,5). RP has been associated with mutations in more than 90 genes – most of which are involved in the function and structure of retinal cells (omim.org/entry/268000#inheritance (3,6)). However, mutations affecting proteins involved in the function and regulation of the spliceosome are collectively the second most common cause of autosomal dominant RP (adRP) (2,7). The spliceosome is a large ribonucleoprotein complex that assembles from its building blocks – the small nuclear ribonucleoproteins (snRNPs) named U1, U2 and U4/U6.U5 (8). Most splicing factor mutations causing RP have been found in three proteins, PRPF31, SNRNP200, and PRPF8 (4), which are all components of the U4/U6.U5 tri-snRNP (4,9,10). PRPF8 is a highly conserved central scaffolding protein of the U4/U6.U5 tri-snRNP and the largest protein in the spliceosome (10–12). It interacts with and regulates the RNA helicase SNRNP200 that unwinds the U4/U6 RNA duplex during spliceosome activation (13,14). Interestingly, the RNA helicase SNRNP200 is also a target of RP-associated mutations, which affect binding of U4 and U6 snRNAs, U4/U6 unwinding, and splicing kinetics (15–17). PRPF8 interacts with and regulates SNRNP200 via the C-terminal Jab1/MPN (10,14). RP mutations cluster exclusively in this domain and we have previously shown that most PRPF8 mutations prevent the efficient formation of the U4/U6.U5 tri-snRNP (10). However, the single substitution p.Tyr2334Asn (Y2334N) at the very C-terminus of PRPF8 does not alter the interaction with SNRNP200 nor prevents the formation of the U4/U6.U5 tri-snRNP (10). In general, it remains elusive how the Y2334N mutation and other mutations in splicing factors that do not disrupt snRNP biogenesis lead to retina-restricted defects.

Photoreceptors are structurally divided into three major parts: the cell body, the photoreceptor inner segment, and the photoreceptor outer segment (POS). The POS is constantly renewed, while the distal part of the POS is shed and phagocytosed by RPE multiple times throughout the day to recycle the used photopigments (6,18,19). Disruptions in the POS renewal process and stability have been shown to manifest in various forms of RP, but due to the interplay between photoreceptors and RPE, it has been difficult to determine whether the RP variants primarily affect photoreceptors or RPE (6,19,20). Various studies of RP-associated mutations in pre-mRNA processing factors in mouse models identified morphological and functional defects in the RPE. At the same time, photoreceptors remained largely unaffected, leading to the paradigm that RPE cells are the primary affected cell type in the retina, followed by photoreceptor loss due to RPE atrophy (21–24). Recently, we generated a PRPF8^Y2334N/Y2334N^ mouse strain that did not show any photoreceptor defects or visual malfunction, seemingly corroborating these findings (25). However, we observed severe cerebellar atrophy and circular RNA expression dysregulation (25). circRNAs are a special class of RNA molecules created from pre-mRNAs by back-splicing when a 5’ splice donor site of an exon is joined to the 3’ splice acceptor site of an upstream exon, resulting in a circular transcript. Due to the lack of 5’ and 3’ ends, these RNAs are remarkably stable and often expressed in neurons and retinal cells (26–29). In addition, differential circRNA expression has been associated with human diseases such as cancer, neurological disorders, and retinal dystrophies, but the molecular mechanism remains mainly unclear (27,30–34).

Advancements in stem cell differentiation techniques have transformed retinal disease modeling by enabling the generation of human induced pluripotent stem cells (hiPSCs)-derived *in vitro* model systems. Recent attempts to extend the mouse model studies of PRPF8 mutations to human retinal cells by employing RP patient hiPSC-derived RPE cells did not reveal any major morphological or functional defects in RPE, challenging the paradigm that RPE are the primary affected cell type, at least in humans (35–37). Here, we combined CRISPR/Cas9 genome editing with an established retinal differentiation protocol to study how the Y2334N variant in PRPF8 affects the neural retina independently of the RPE. We recapitulated photoreceptor outer segment defects in retinal organoids harboring the Y2334N mutation, revealing a disease-relevant phenotype in human photoreceptors. Using transcriptome analysis, we found that the Y2334N mutation induces splicing alterations in a subset of genes with retinal function and dysregulation of specific circular RNAs.

## Materials and Methods

### iPSC maintenance

Human induced pluripotent stem cells (hiPSC) were generated and characterized as described in (38). hiPSCs were cultured in six-well plates on wells coated with recombinant, truncated human vitronectin (Thermo Fisher Scientific, A14700) in Essential 8 medium or Essential 8 Flex medium (Thermo Fisher Scientific, A2858501). According to the manufacturer’s instructions, the cell culture medium was replaced daily or thrice a week. Cells were passaged every 3-4 days by incubation in 0.5 mM EDTA/PBS at 37 °C for 3-5 minutes and transferred to vitronectin-coated wells in a 1:5 – 1:10 ratio. hiPSCs were maintained at 37 °C and 5% CO_2_.

### CRISPR Cas9 genome editing of hiPSC

To introduce the Y2334N mutation in *PRPF8* and a C-terminal GFP tag (or to add the GFP tag only), we used CRISPR-Cas9 genome editing as previously described (25). In brief: The guide RNA sequence was designed using the CRISPR design tool (39). The guide RNA was cloned into the pX330-U6-Chimeric BB-CBh-hSpCas9 plasmid (# 42230; Addgene). The homology-directed repair (HDR) template was assembled from 1.5kb left and right homology arms flanking the target sequence and a GFP sequence with SV40pA. The homology arms were amplified by PCR from genomic DNA, and the target sequences with GFP were amplified from PRPF8-pEGFP-N1 and Y2334N-pEGFP-N1 plasmids. hiPSCs were grown to 80% confluence and co-transfected with linearized HDR template and pX330 plasmid harboring the guide RNA sequence using Lipofectamine Stem Transfection Reagent (Thermo Fisher Scientific) according to the manufacturer’s protocol. Non-homologous end-joining inhibitor SCR7 (1 µM) was added 24 hours before transfection and replenished every other day until fluorescence-activated cell sorting (FACS). GFP-positive cells were sorted 72 hours post-transfection. Positive clones were selected by PCR and Sanger sequencing to confirm correct editing. The top five predicted off-target sites were verified as unmodified.

### Immunoprecipitation

Cells grown on 100 mm diameter Petri dishes were washed with ice-cold PBS, harvested by centrifugation, resuspended in NET2 buffer (50 mM Tris-HCl pH 7.5, 150 mM NaCl, 0.05% Nonidet P-40) supplemented with protease inhibitor cocktail, and pulse sonicated for 60 s on crushed ice. The cell lysate was centrifuged and the supernatant was immunoprecipitated with goat anti-GFP antibodies (kindly provided by E. Geertsma, MPI-CBG, Dresden, Germany) bound to Protein G PLUS – Agarose beads (Santa Cruz Biotechnology). The immunoprecipitated proteins were separated on a 4-15% gradient SDS-PAGE, transferred to nitrocellulose membrane and individual proteins immunodetected.

### Western blotting and immunodetection

proteins were separated on a 4-15% gradient SDS-PAGE, transferred to nitrocellulose membrane and immunodetected using the following antibodies: anti-PRPF8 (AB79237, Lot GR3216012-3), anti-EFTUD2 (AB72456) and anti-PRPF31 (AB188577) from Abcam, anti-SNRNP200 (Sigma-Aldrich, HPA029321, Lot 000049736), anti-PRPF6 (Santa Cruz Biotechnology, sc166889, Lot 1713), anti-GFP (Santa Cruz Biotechnology, sc9996, Lot#: E1123), anti-PRPF4 (kindly provided by R. Lührmann, MPI, Göttingen, Germany), anti-tubulin antibody used as a loading control (kindly provided by P. Dráber, IMG, Prague, Czech Republic). Secondary anti-mouse and/or anti-rabbit antibodies conjugated with horseradish peroxidase were used for western blotting detection (Jackson ImmunoResearch Laboratories).

### Immunofluorescence

Cells were grown on vitronectin-coated coverslips, washed with PBS, fixed with 4% paraformaldehyde/PIPES for 10 min at RT and permeabilized with 0.2% (v/v) Triton X-100 in PBS containing 5% normal goat serum (NGS). Cells were incubated with primary mouse anti-SRSF2 antibody (kindly provided by K. Neugebauer, Yale, New Haven), followed by incubation with secondary anti-mouse DyLight 549 antibody (Jackson ImmunoReseach Laboratories). Stained cells were dried out and mounted with Fluoromount G (Southern Biotech) containing DAPI for DNA visualization. Images were acquired with an Olympus IX83 equipped with an oil immersion 60x objective (1.42 NA).

### hiPSC differentiation to retinal organoids

hiPSCs differentiation into retinal organoids was performed based on a previously published protocol (40) with minor modifications. Briefly, hiPSCs were dissociated into single cells using EDTA/PBS and 3000 cells were aggregated in low-adhesion 96-well plates in Essential 8 medium (Thermo Fisher Scientific, A2858501) with 20 µM ROCK inhibitor Y-27632 (Tocris, 1254). 48h later (defined as day 0) the medium was changed to retinal differentiation medium (RDM) composed of 5% Iscove’s modified Dulbecco’s medium (IMDM, 12440046; Thermo Fisher Scientific), 45% Ham’s F12 (Thermo Fisher Scientific, 11765054), 10% KnockOut Serum Replacement (KSR; Thermo Fisher Scientific, 10828028), GlutaMAX (Thermo Fisher Scientific, 35050038), 1% Chemically Defined Lipid Concentrate (Thermo Fisher Scientific, 11905031), 450 µM monothioglycerol (Sigma-Aldrich, M6145), and penicillin/streptomycin (Thermo Fisher Scientific, 15140–122). On day 6, BMP4 (Peprotech 120-05ET, Lot#: 0419526, Lot#: 0421526) was added to each well at 55 ng/ml. Every third day until day 18, half of the medium was replaced with fresh RDM. On day 18, retinal organoids were transferred to Petri dishes and cultured in organoid long-term (OLT) medium composed of DMEM/F12 (Thermo Fisher Scientific, 31330–038), 10% fetal bovine serum (FBS; Thermo Fisher Scientific, 10270–106), 1% N2 Supplement (Thermo Fisher Scientific, A1370701), 0.1 mM taurine (Sigma-Aldrich, T8691), 0.5 µM retinoic acid (Sigma-Aldrich, R2625), 0.25 µg/ml Amphotericin B (Thermo Fisher Scientific, 15290-02), and penicillin/streptomycin (Thermo Fisher Scientific, 15140-122). Retinal organoids were maintained in OLT for up to 250 days with medium changes every 3-4 days.

### Immunohistochemistry staining of retinal organoids

At day 170, retinal organoids were transferred to a microcentrifuge tube and rinsed with PBS. Organoids were fixed in 4% paraformaldehyde solution for 30 minutes, followed by overnight incubation in 30% sucrose in PBS at 4 °C. Sucrose was removed, and organoids were embedded in OCT (Sakura Finetek, 4583) and 10 µm sections were prepared on a Cryostat Leica CM1950. Sections were permeabilized and blocked using 0.3% Triton-X-100, 10% normal goat serum (SouthernBiotech, 0060-01) in PBS. Sections were stained for retinal marker proteins overnight at 4 °C with the following primary antibodies: RHODOPSIN (Sigma-Aldrich, O4886, Lot#: 22180131, 1:200), RECOVERIN (Sigma-Aldrich, AB5585, Lot#: 3432603, 1:1000), CRALBP (Abcam, ab15051, Lot#: GR3284867-10 1:200), and AP-2α (Santa Cruz, sc-12726 Lot#: E1921, 1:100). The following secondary antibodies were used: goat anti-mouse IgG Alexa Fluor^TM^ 555 (Thermo Fisher Scientific, A21424, Lot#: 1761228, 1:750), goat anti-rabbit IgG Alexa Fluor^TM^ 555 (Thermo Fisher Scientific, A21429, Lot#: 1750835, 1:750), goat anti-mouse IgG Alexa Fluor^TM^ 647 (Thermo Fisher Scientific, A21236, Lot# 2119153, 1:750), and goat anti-rabbit IgG Alexa Fluor^TM^ 647 (Thermo Fisher Scientific, A21245, Lot#: 2299231, 1:750). Nuclei were counterstained with DAPI (Sigma-Aldrich, 10236276001), samples were mounted using VECTASHIELD HardSet (Vector Laboratories, H1400) and images were acquired on a Dragonfly 503 spinning disk confocal microscope (Andor) equipped with an oil immersion HCX PL APO 40x objective (1.25 NA) and processed using Fiji (ImageJ).

### Transmission electron microscopy

Retinal organoids were fixed using 3% glutaraldehyde (Sigma Aldrich, G5882) in 0.1 M sodium cacodylate buffer (pH 7.3, Sigma Aldrich, C0250) at 4°C. After washing with 0.1M sodium cacodylate buffer, samples were post-fixed in 1% OsO4 (Sigma Adrich, 05500) in 0.1M sodium cacodylate buffer at room temperature for 1.5h. After another washing step in 0.1M sodium cacodylate buffer, the samples were dehydrated in ethanol series (50%, 70%, 96% and absolute ethanol, Penta, #71250-11002) and acetone and embedded in the DurcupanTM ACM resin (Sigma Aldrich, resin composed from four components: A44611+B44612+C44613+D44614). During a three-day resin polymerization process, the temperature gradually increased from 60°C to 80°C. The ultrathin sections of the samples were cut using an ultramicrotome Leica EM UC6 (Leica Mikrosysteme GmbH) and examined using a Morgagni 268D transmission electron microscope (Thermo Fisher Scientific, Eindhoven, Netherlands), working at 90 kV and equipped with a Veleta CCD camera (Olympus, Münster, Germany).

### Scanning electron microscopy

Retinal organoids were fixed using 3% glutaraldehyde (Sigma Aldrich, #G5882) in 0.1 M sodium cacodylate buffer (pH 7.3, Sigma Aldrich, #C0250) at 4°C. After fixation, the samples were gently rinsed with 0.1 M sodium cacodylate buffer and subjected to dehydration through a graded series of ethanol solutions (30%, 50%, 70%, 80%, 90%, 96%, and absolute ethanol) (Penta, #71250-11002). Following dehydration, the samples were critical point-dried (CPD 030, BAL-TEC Inc.) using liquid CO_2_. Once dried, the samples were coated under vacuum with a thin layer of gold using a Bal-Tec SCD 040 sputter coater (LEICA, Wetzlar, Germany). The gold-coated organoids were then observed under a scanning electron microscope (VEGA TS 5136 XM, Tescan Orsay Holding).

### Light microscopy analysis and quantification of photoreceptor inner/outer segment layer thickness

Brightfield images were acquired on a Leica DMi8 equipped with an HC PL FLUOTAR L 20x/0.40 DRY objective. The brush borders were segmented using the *Trainable WEKA Segmentation* plugin available in Fiji (ImageJ). The underlying machine learning algorithm was trained on representative images of PRPF8-WT and PRPF8-Y2334N retinal organoids containing POS structures. Once trained, the model was applied to brush border regions in 12 images of three biological replicates of PRPF8-WT and PRPF8-Y2334N retinal organoids. Where necessary, manual corrections were made on the segmented images to eliminate artifacts and ensure accurate delineation of the inner/outer segment region before quantitative analysis. Following segmentation and manual refinement, a custom macro written in the ImageJ Macro Language (IJM) was used to automatically quantify the average thickness of the inner/outer segment regions. The analysis combined two Fiji plugins:

*Skeletonize (2D/3D)* and *Analyze Skeleton (2D/3D)*: These tools reduced the segmented areas to their central lines (skeletons) and computed the total length of each skeletonized structure.

*Local Thickness*: This plugin calculates the local thickness at each point within the segmented POS regions.

The macro integrates the outputs of these tools to evaluate both the length and the average thickness of each POS structure. The analysis yields the following outputs:

– A labeled skeleton image.
– A local thickness map.
– A CSV file containing quantitative statistics for each skeleton, including its length and average thickness.

### RNA isolation, library preparation and RNA-seq

Retinal organoids were rinsed with PBS, immediately collected in TRIzol reagent (Thermo Fisher Scientific, 15596018) and stored at −80 °C. Retinal organoids were lysed in TRIzol reagent and total RNA was extracted using the Direct-Zol RNA Microprep Kit (Zymo Research, R2062), including on-column DNase-I treatment. The quantity and quality of isolated RNA were measured using NanoDrop ND-2000 (NanoDrop Technologies) and analyzed by Agilent 2100 Bioanalyser (Agilent Technologies). Depletion of rRNA was done with KAPA RiboErase Kit (HMR) and sequencing libraries were generated using KAPA RNA HyperPrep Kit. Libraries were sequenced on the Illumina NextSeq® 500 instrument using a 150 nt paired-end configuration (2x 75 nt).

### Analysis of alternative splicing and circular RNA expression

Sequencing reads were processed using the bioinformatic pipeline nf-core/rnaseq v1.4.2 (41,42). Individual steps included removing sequencing adaptors and low-quality reads with Trim Galore v0.6.4 (43) and cutadapt v2.5 (44), mapping to reference genome GRCh38 (Ensembl annotation version 101) (45) with HISAT2 v2.1.0 (46). Bam files generated by HISAT2 were processed using ASpli v2.2.0 (46) R Bioconductor package to conduct gene-level differential expression analysis and quantify splicing efficiency for annotated alternative splicing exons and introns. This included differential analysis based on exon exclusion/intron retention supporting reads and calculating the difference of exclusion (PSI) and retention (PIR) ratios between Y2334N and WT groups. Furthermore, perturbation of circular RNA (circRNA) profiles between Y2334N and WT groups was assessed based on back-spliced reads using CIRI v2.0.6 (47) for circRNAs quantification from raw reads and CIRIquant v1.1.1 (48) for differential expression analysis of individual circRNAs.

### RT-qPCR for quantification of differentially expressed circRNAs

Retinal organoids were rinsed with PBS, immediately collected in TRIzol reagent (Thermo Fisher Scientific, 15596018) and stored at −80 °C. Retinal organoids were lysed in TRIzol reagent and total RNA was extracted using the Direct-Zol RNA Microprep Kit (Zymo Research, R2062), including on-column DNase-I treatment. 250 ng of total RNA was reverse transcribed using SuperScript III Reverse transcriptase (Invitrogen) and random hexamers (Invitrogen). The cDNA obtained was diluted 1:10 in a 5 μL reaction for qPCR by LightCycler480 (Roche) to measure the extent of circRNA and linear mRNA biogenesis. The first set of primers was designed in reverse orientation to span the back-spliced junction between the 5’ end of the first exon and the 3’ end of the second exon, which determines the circRNA. The second set of primers faces each other, spanning the junction between the 3’ end of the second exon and the 5’ end of the third exon. This amplifies the linear spliced variant. The fold change expression level (2-ΔΔCt) was measured as a circular/ circular+linear RNA ratio between the wild type and mutant organoids. Three reference genes-GAPDH, ACTB, and H2B were used as internal controls.

The following primers were used:

1. PB454_cSTK39_F: CCATCTCGTCGTCACTCCAC
2. PB455_cSTK39_R: AGCTTCTTCTTGTGCCGTGA
3. PB456_linSTK39_R: ACCAGACATAGCCCAAAGAGC
4. PB281_circPDK1_F: TGAAAATGCTAGGCGTCTGTG
5. PB282_circPDK1_R: CATCCTCAGCACTTTTGTCCTT
6. PB283_linPDK1_R: CACTTGTATTGGCTGTCCTGG

### Analysis of splice site strength

Splice site analysis was performed using the human GRCh38 genome assembly as the reference, with intron coordinates obtained from GENCODE (version 47) (49). For each intron, sequences spanning positions −3 to +6 relative to the 5′ splice site (5′ss) and positions −20 to +3 relative to the 3′ splice site (3′ss) were extracted using a custom Python script. Sequence retrieval was carried out using BEDTools getfasta (v2.30.0) (50). The strength of all 5′ss and 3′ss was calculated using the Maximum Entropy (MaxEnt) algorithm (51). Splice site scores were computed individually for differentially more and differentially less retained introns in the Y2334N mutant. All statistical analyses and visualizations were performed in R (version 4.0.4), with boxplots generated using the ggplot2 package.

## Results

### Generation and characterization of hiPSC-derived retinal organoids

We applied CRISPR/Cas9 genome editing to introduce the p.Tyr2334Asn substitution together with a C-terminal GFP tag into one allele of the *PRPF8* gene in hiPSC derived from peripheral blood mononuclear cells of a healthy donor. At the same time, we GFP-tagged a wild-type allele of the *PRPF8* gene as a control to exclude effects of the GFP tag. The insertion of the mutation and the tag was verified by genome sequencing (Fig. S1A). The tagged PRPF8 was expressed at the same level as the untagged protein (Fig. S1B). Surprisingly, both endogenous and the tag variant expression were lower in hiPSC carrying the Y2334N mutation, indicating that the mutation reduces the PRPF8 protein levels. Furthermore, neither the GFP tag at the C-terminus alone nor in combination with the Y2334N mutation affected the correct localization of the PRPF8 protein (Fig. S1C). Hereafter, we refer to the hiPSC line with the p.Tyr2334Asn substitution as “PRPF8-Y2334N” and the hiPSC line without the mutation as “PRPF8-WT”.

It has been previously published that in HeLa cells, the Y2334N mutation does not affect the interaction of PRPF8 with SNRNP200 and other tri-snRNP-specific proteins (10). To confirm that this also applies to hiPSCs, we immunoprecipitated PRPF8-Y2334N-GFP and PRPF8-WT-GFP using anti-GFP antibodies. Our results are consistent with previous findings and show that the Y2334N mutation does not interfere with the interaction of PRPF8 with SNRNP200 and other tested snRNP-specific proteins (Fig. 1).

**Figure 1.**
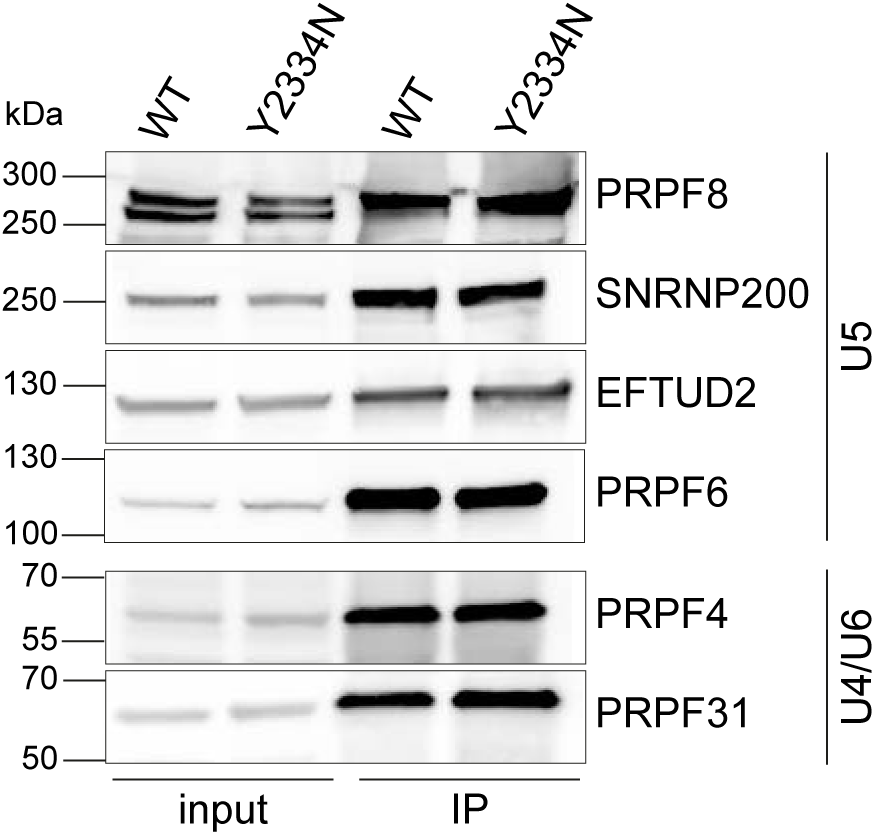
The Y2334N mutation does not impair the interaction of PRPF8-Y2334N with U5 and U4/U6.U5 snRNP proteins in hiPSC. PRPF8-GFP was immunoprecipitated by anti-GFP antibodies and specific proteins were detected by western blotting. Inputs contained 2% of the total lysate.

After we confirmed that this unique property of the Y2334N variant is preserved in hiPSC, we differentiated PRPF8-Y2334N and PRPF8-WT hiPSCs into retinal organoids. The Y2334N mutation did not affect the early differentiation potential of the PRPF8-Y2334N cell line and both PRPF8-WT and PRPF8-Y2334N developed an emerging neural retina after 18 days (Fig. 2A). At 170 days after initiation of differentiation, retinal organoids derived from both PRPF8-WT and PRPF8-Y2334N hiPSCs possessed a structured and laminated neural retina that contained photoreceptors (stained by recoverin/rhodopsin, Müller (stained by CRALBP) and amacrine (stained by AP-2α) cells (Fig. 2B). We observed some differences in retinal organization of WT and Y2334N organoids, but these differences are rather caused by the variability of individual organoids than the Y2334N allele. We concluded that the Y2334N mutation did not affect early retinal development and the overall cellular composition and organization of retinal organoids at an early stage. This finding is consistent with the clinical picture of RP, characterized by a late disease onset and postnatal initiation of cerebral degeneration in our mouse model (25).

**Figure 2.**
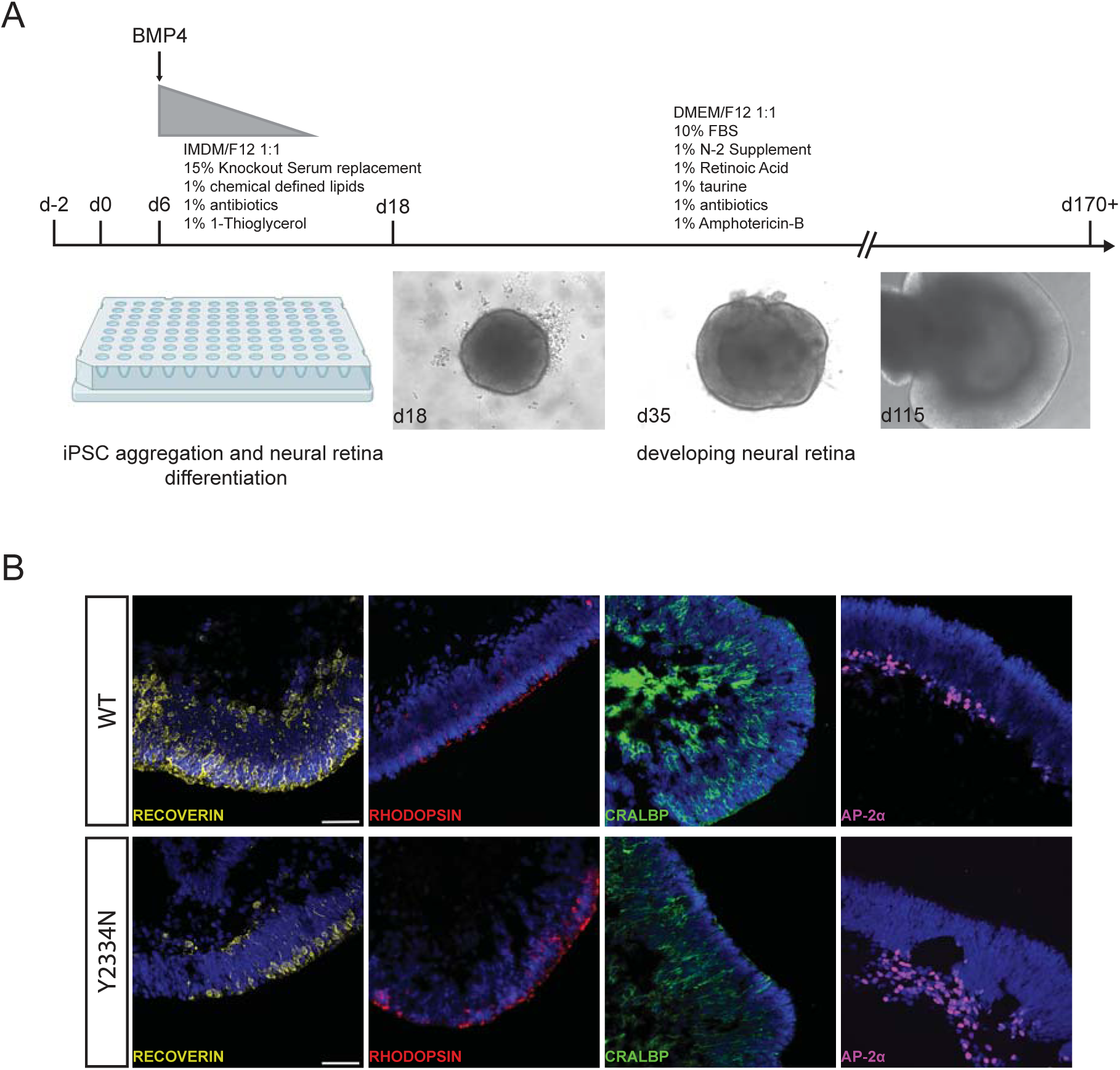
Generation of 3D retinal organoids from PRPF8-WT and PRPF8-Y2334N hiPSC. A) Schematic representation of the differentiation of hiPSC to 3D retinal organoids. Representative brightfield images illustrating the development of the neural retina during differentiation. Created in BioRender. Stanek, D. (2025) https://BioRender.com/2aaivccB) Comparative immunohistochemistry staining of 170-day-old retinal organoids, derived from PRPF8-WT and PRPF8-Y2334N hiPSC, showing the expression of indicated retinal markers. DNA was stained by DAPI (blue). Scale bars - 50 μm.

### Retinal organoids differentiated from PRPF8-Y2334N hiPSC show a lower density of photoreceptor outer segments

We then directed our attention to photoreceptors to assess the potential of the Y2334N mutation to alter their morphology, particularly the POS. We cultivated retinal organoids for 250 days and analyzed POS using electron microscopy. First, we applied transmission electron microscopy but found no significant changes in the ultrastructure of the photoreceptor cell body and inner segment (Fig. 3). The POSs were rarely present, and we observed them only in PRPF8-WT organoids. Next, we monitored the surface of retinal organoids by scanning electron microscopy. The surface of organoids derived from PRPF8-WT hiPSCs was entirely covered with structures resembling inner and/or outer segments and we call them inner-like segment structures. In contrast, PRPF8-Y2334N retinal organoids revealed only a sparse coverage of the organoid’s surface with these inner-like segment structures. This indicates that retinal photoreceptors expressing the RP mutant fail to develop and/or maintain outer segments, and that the density of inner segments is altered in organoids expressing the pathogenic variant of PRPF8. However, it should be noted that outer segments are often lost during sample preparation for electron microscopy (52).

**Figure 3.**
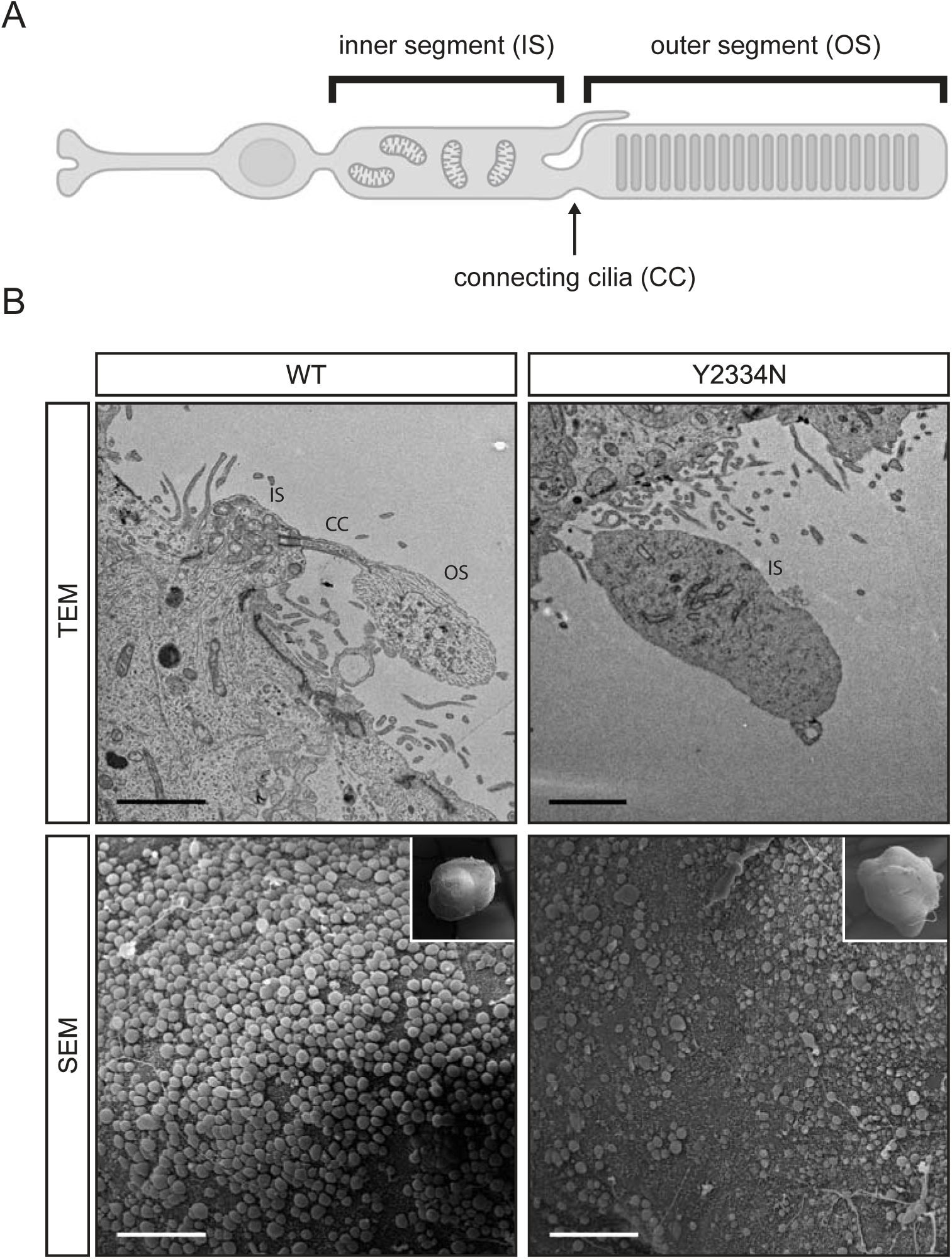
Y2334N retinal organoids show impaired photoreceptor inner segments. A) Schematic representation of the photoreceptor structure. Created in BioRender. Stanek, D. (2025) https://BioRender.com/8bgoqrx. B) Transmission electron microscopy (TEM) images of photoreceptors from 250-day-old retinal organoids. Outer segments were visible in the WT but not the Y2334N retinal organoids (upper panel). Scanning electron microscopy (SEM) images of 250-day-old retinal organoids revealed fewer inner segments on the retinal organoid surface of Y2334N organoids (lower panel). Scale bars: Top panel - 2 μm, bottom panel - 50 μm. IS = inner segment, CC = connecting cilia, OS = outer segment.

To overcome the loss of POS during sample preparation for electron microscopy or immunohistochemistry, we performed live cell imaging of retinal organoids (Fig. 4A). We observed that the layer above the outer nuclear layer, containing inner and outer segments called the brush border, is thinner in PRPF8-Y2334N organoids. We employed a machine learning-supported image segmentation analysis to quantify the thickness of the brush border. We observed a 50% reduction in the average the brush border thickness of PRPF8-Y2334N retinal organoids compared to PRPF8-WT organoids (Fig. 4B). In addition, the area covered by photoreceptor segments seemed smaller in organoids derived from PRPF8-Y2334N hiPSCs, which was consistent with scanning electron microscopy results (Fig. 3B). However, the difference did not pass the statistical significance test (Fig. 4C). Taken together the Y2334N mutation does not affect the inner organization of photoreceptors but prevents the correct formation of outer segments.

**Figure 4.**
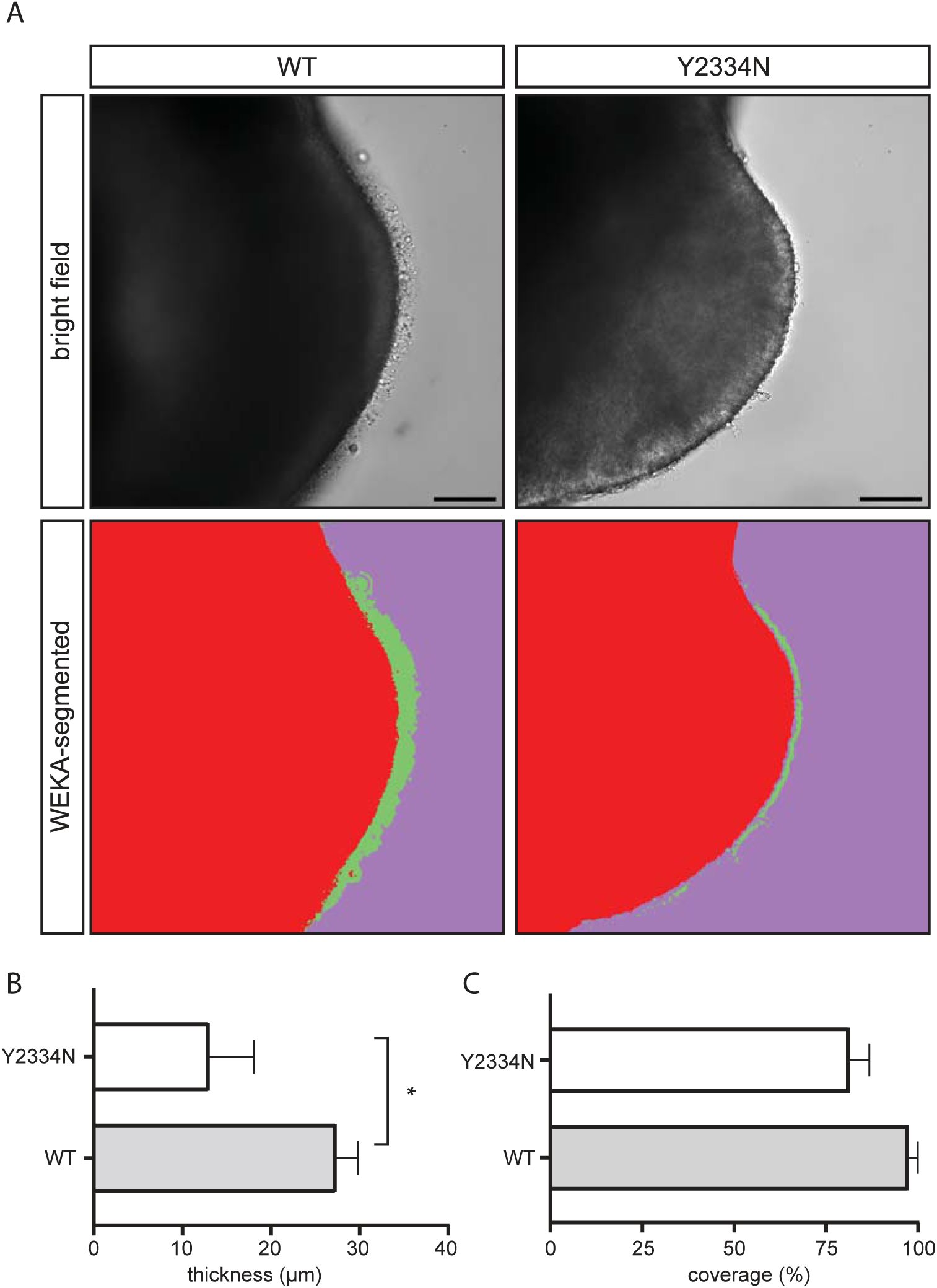
The thickness of the photoreceptor brush border is reduced in Y2334N retinal organoids. A) Bright field images of a representative neural retina from 250-day-old retinal organoids (upper panel). Machine-learning-supported WEKA segmentation analysis of the respective brightfield image (lower panel). The photoreceptor inner and outer segment layers are depicted in green. Scale bar 50 μm, B) Quantification of the photoreceptor inner and outer segment layer thickness. The layer is significantly thinner in Y2334N retinal organoids compared to WT organoids. C) Quantification of inner segment coverage on the organoid surface. No significant difference in the relative coverage was detected. Data are represented as mean +/- SEM of three biological replicates. Statistical significance was determined using a two-tailed unpaired t-test (* indicates p < 0.05).

### Transcriptome analysis reveals differential splicing of retinal genes

We performed transcriptome analysis to get better insight into molecular defects in Y2334N organoids and how this might translate into POS malformation. The transcriptome of Y2334N and WT organoids was similar, with only two genes, TACR3 and SLITRK2, being significantly upregulated in Y2334N organoids (Fig. 5A, Suppl. table 1). SLITRK2 is a gene previously associated with synapse formation and neurite outgrowth. Analysis of existing single-cell RNA-seq data of the human retina (53) revealed that SLITRK2 is predominantly expressed in Müller cells and TACR3 is predominantly expressed in bipolar cells (Fig. 5B).

**Figure 5.**
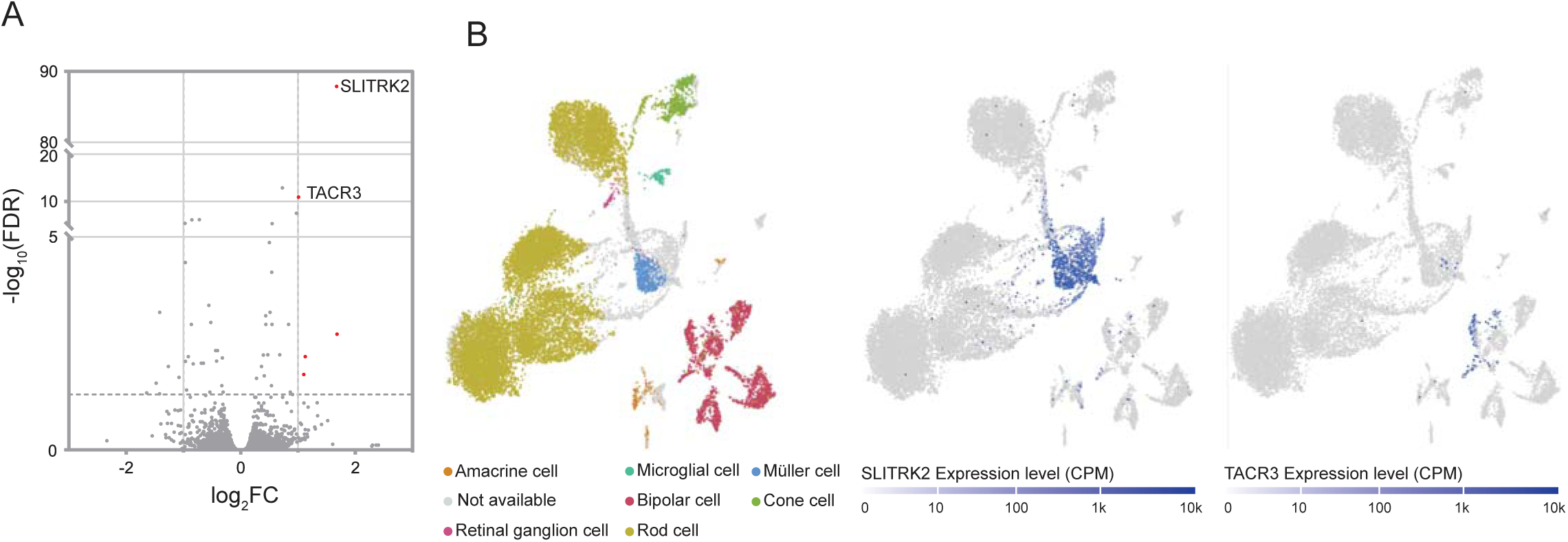
170-day-old PRPF8-Y2334N retinal organoids reveal limited changes in gene. A) Differential expression analysis of PRPF8-WT vs. PRPF8-Y2334N retinal organoids. Genes were considered differentially expressed if FDR < 0.05 and fold change expression ≥ 2 (|log2 FC| ≥ 1). B) Analysis of publicly available single-cell RNA sequencing data from the Single Cell Expression Atlas (E-MTAB-7316) revealed predominant SLITRK2 expression in Müller cells and TACR3 in bipolar cells.

Subsequently, we analyzed whether the Y2334N mutation affects RNA splicing. Initially, we focused on intron retention and found that 568 introns were differentially retained (Suppl. table 2). Compared to WT retinal organoids, Y2334N retinal organoids exhibited increased retention of 303 introns, while 265 introns showed higher retention in WT organoids (Fig. 6A). Strikingly, the intron with the highest differential splicing was found in the *A2M* gene. This intron was 36% more retained in Y2334N retinal organoids compared to WT organoids (Fig. 6A, B). The *A2M* gene has been previously linked to the Müller cell-mediated retinal stress response (54–56) and associated with neural diseases such as Alzheimer’s (57). To better understand why retained introns are susceptible to retention, we compared the splice site strength of the differentially retained introns with other annotated introns of the human genome. We observed that introns with higher retention in the Y2334N retinal organoids possess significantly weaker average 3’ splice sites (Fig. S2A). We observed a similar tendency for the 5’ splice sites, but here the difference was smaller and not statistically significant (Fig. S2B).

**Figure 6.**
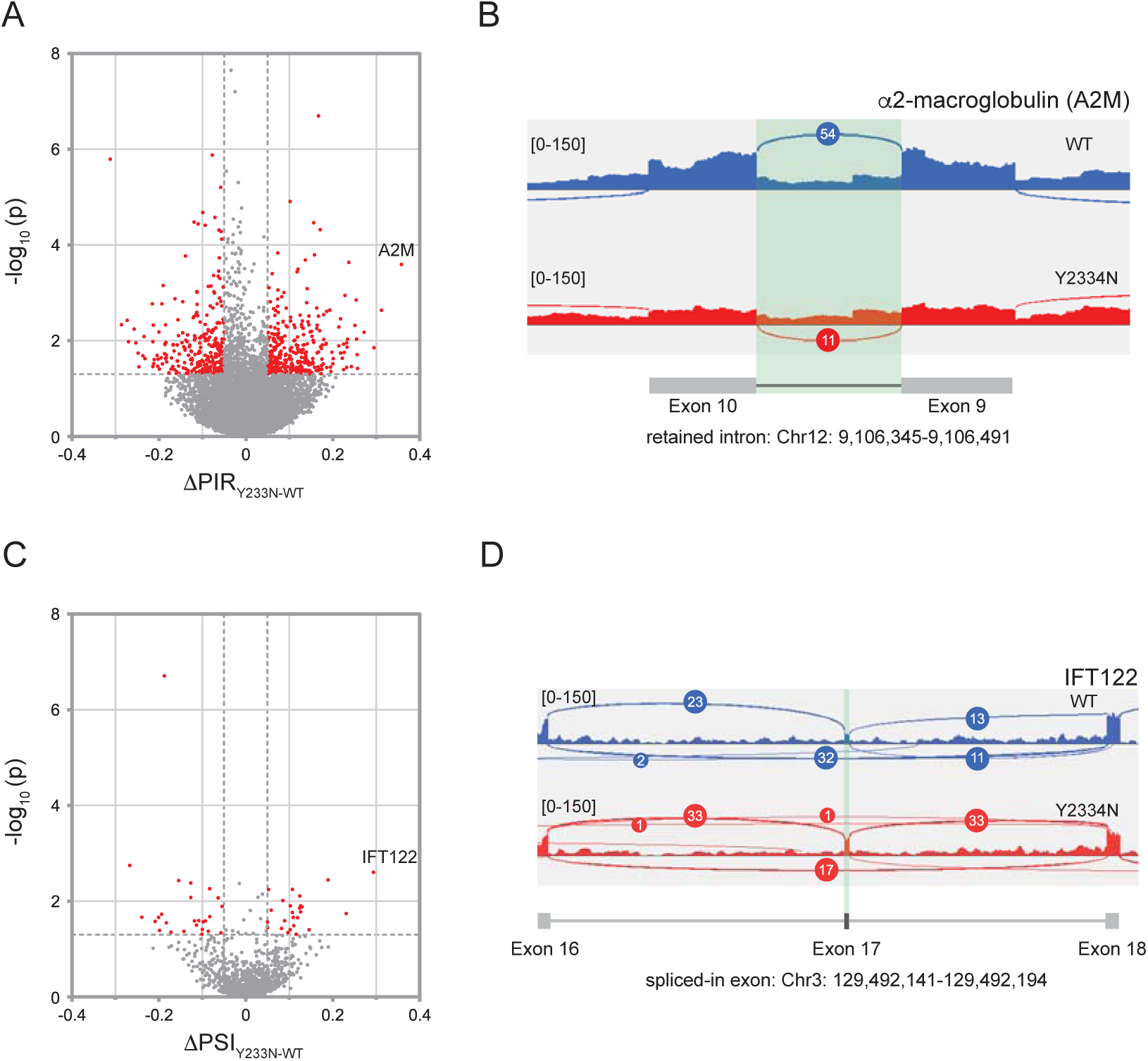
The Y2334N mutation causes changes in splicing in 170-day-old retinal organoids. A) Differential intron retention analysis of Y2334N and WT retinal organoids. Introns were considered differentially retained if ΔPIR^Y2334N-WT^ (difference in percentage intron retention) ≥ 0.05 and p < 0.05. B) The sashimi plot shows the indicated transcript and illustrates the intron with the largest difference in intron retention. The retained intron is highlighted in green. C) Differential analysis of alternative exons (exon skipping). Transcripts were considered differentially spliced if ΔPSI^Y2334N-WT^ (percentage spliced in) ≥ 0.05 and p < 0.05. D) The sashimi plot shows the indicated transcript and illustrates the exon with the most substantial difference in alternative splicing. The alternatively spliced exon is highlighted in green. Each Sashimi plot was generated using data from three merged samples.

We then analyzed alternative splicing and detected 50 alternative exons to be differentially spliced (Fig. 6C, Suppl. table 3). 24 exons were skipped more often in Y2334N retinal organoids and 26 more often in WT organoids. Interestingly, the exon with the highest Percentage Spliced In (PSI) value of 30% was found in the IFT122 gene (Fig. 6D). The IFT complex is required for ciliogenesis and ciliary protein trafficking and is crucial for the function of photoreceptors (58–61). Moreover, dysfunctional IFT proteins were described as a common cause of RP and other progressive retinopathies (58,59,62–64).

These data indicate that the Y2334N mutation causes rather limited changes in the RNA landscape and alters splicing of a specific subset of retinal transcripts, particularly IFT proteins, that may be critical drivers of the RP phenotype in human retinal cells. Interestingly, our data suggest that the Y2334N PRPF8 mutation not only affects photoreceptors, but molecular markers of cellular stress were also detected in other cells of the retina, e.g., Müller cells.

### Circular RNAs are dysregulated in PRPF8-Y2334N retinal organoids

We recently discovered that the Y2334N mutation correlates with differentially expressed circRNAs in the cerebellum of mice expressing Prpf8-Y2334N (25). Intending to extend our circRNA studies from the mouse cerebellum to the human neural retina, we investigated whether circRNAs are also dysregulated in the human retinal organoid model of RP. We prepared an unbiased sequencing library without selection of poly-adenylated transcripts from total RNA isolated from PRPF8-Y2334N and PRPF8-WT retinal organoids 170 days after the start of differentiation. Our analysis revealed 102 circRNAs exhibiting differential expression in PRPF8-Y2334N, with 59 circRNAs showing upregulation and 43 circRNAs showing downregulation (Fig. 7B, Suppl. table 4). To get a better understanding of how circRNA expression develops in aging organoids, we selected two circRNAs, PDK1_hsa_circ_0141757 and STK39_hsa_circ_0003279, and monitored their expression over time using RT-qPCR (Fig. 7C). We observed an age-dependent accumulation of these circRNAs in retinal organoids, and this accumulation was elevated in Y2334N organoids compared to WT organoids. Given the age-dependent increase in circRNA expression in retinal organoids, circRNAs could serve as potential biomarkers of retinal atrophy progression, particularly in Y2334N-mediated RP.

**Figure 7.**
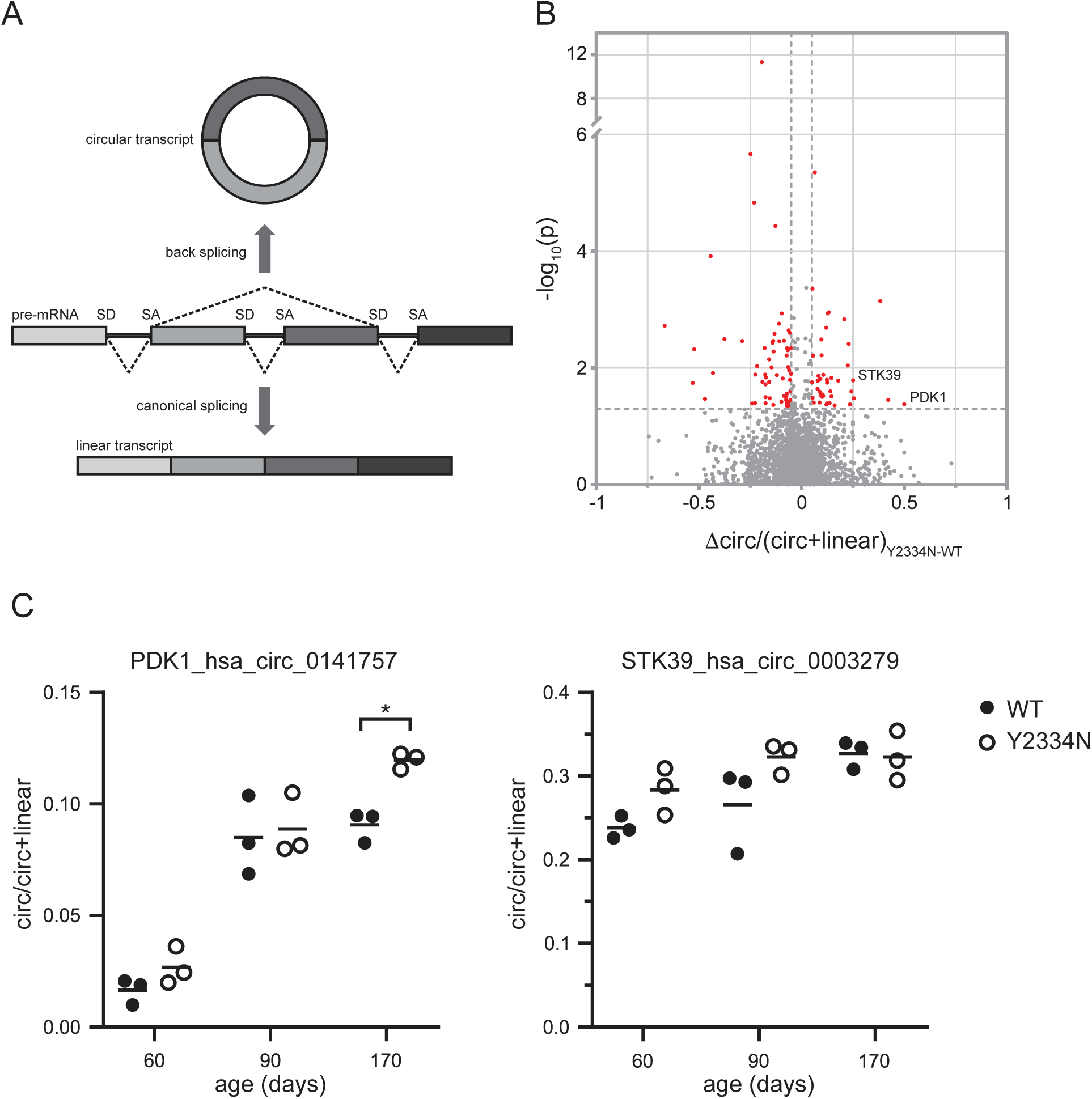
Circular RNA expression is changed in PRPF8 Y2334N retinal organoids. A) Schematic representation of circRNA splicing-dependent production. B) Differential expression of circRNAs between WT and Y2334N retinal organoids. circRNAs were considered differentially expressed if Δratio(circ/circ+lin)^Y2334N-WT^ ≥0.05 and p < 0.05. C) Developmental expression profiles of PDK1_hsa_circ_0141757 and STK39_hsa_circ_0003279 circRNAs in retinal organoids, as determined by RT-qPCR. Data are represented as the mean of three biological replicates. Statistical significance was determined using a two-tailed unpaired t-test (* = p < 0.05).

## Discussion

Mutations in splicing factors represent the second most common cause for adRP. In many cases, the tri-snRNP proteins PRPF31, PRPF8 and SNRNP200 are affected. The mechanistic effects of these mutations on the spliceosome and splicing have been extensively studied in immortalized cell lines and mouse models. Notably, insights from mouse models established the paradigm that cells of the RPE are the primary affected cell type. However, recent studies on human RPE cells differentiated from RP patients challenged this prevailing model, revealing limited/no RPE defects associated with PRPF8 pathological variants (35–37). At the same time, only a few recent studies explored the effect of the PRPF mutations in biological models where the effects of the pathogenic variant on the human neural retina, and specifically on the human photoreceptors, can be mainly evaluated independently of the RPE (36,65,66).

Here, we focused on the Y2334N mutation in the PRPF8 protein and developed a stem-cell-derived retinal organoid model to study the effect on the neural retina. While the overall composition and organization of the neural retina were unaffected by the mutation, we observed reduced thickness of the photoreceptor brush border (Fig. 4). This result is consistent with a recent study showing impaired POS development in retinal organoids carrying the pathogenic p.H2309P variant in one PRPF8 allele (36). This and our findings demonstrate that retinal organoids effectively mimic the RP phenotype, highlighting their utility as a valuable model for dissecting the molecular mechanism underlying RP. In addition, our results suggest that the formation of inner segments on the organoids’ surface might also be compromised (Fig. 3), which indicates that the PRPF8 pathogenic variant impairs the overall stability of both inner and outer segments. In the human retina, photoreceptors directly contact the RPE via the POS. However, the differentiation protocol we employed promotes the development of neural retinal cells, with only minimal RPE presence that remains physically separated from the neural retina (40). Because the interaction between photoreceptor and RPE is not recapitulated in our retinal organoids, pathogenic effects observed in photoreceptors are uncoupled from potential RPE-related defects. Therefore, our findings suggest that photoreceptors are intrinsically affected in RP caused by PRPF8 mutations. This hypothesis is further supported by previous studies reporting a lack of morphological and functional defects in RPE models, alongside a predominant photoreceptor phenotype observed in retinal organoids (35–37).

The predominant manifestation of functional defects in RPE in mouse models may be partially attributed to the substantial differences in photoreceptor and RPE densities between rodent and human retinas (67). In mice, photoreceptor defects, if any, do not show before the mice reach a very high age. In RP mouse models and human RP patients, weakened photoreceptor segments might be stabilized by the retinal environment; therefore, the disease phenotype manifests later in life. *In vitro* retinal organoid models lack this protective retinal environment and photoreceptor segments may disappear much faster, and defects are visible already 30 weeks after the start of differentiation.

Establishing a relevant model for studying adRP allowed us to investigate molecular changes in the RNA landscape that preceded the phenotype in photoreceptors. We compared the transcriptome of WT and Y2334N organoids and found surprisingly few differences. Differential expression analysis revealed significant changes in the expression of only two transcripts, TACR3 and SLITRK2. TACR3 has essential functions in reproduction and the central nervous system (68–70). In the human retina, TACR3 is predominantly expressed in bipolar cells, but its role is largely unknown (71–73). SLITRK2 has been described as a regulator of neurite outgrowth, and upregulation of SLITRK2 has been associated with synaptic dysfunctions (74–76). More importantly, SLITRK2 is predominantly expressed in Müller cells and overexpression of SLITRK2 could indicate defects in this cell type. Müller cells fulfill many functions, from guiding light over structural support to retinal homeostasis and stress response (77–82). In addition, Müller cells are required for the development of photoreceptors and the dynamics of the photoreceptor outer segments (83). This raises the possibility that the deleterious effects of splicing factor mutations are not limited to photoreceptors and the RPE but may impact other neural retina cell types. This conclusion is consistent with our analysis of splicing changes in retinal organoids, which revealed defects in the Alpha-2-macroglobulin (A2M) transcript. A2M is essential for Müller cell-mediated stress response (54,55) and has been previously linked to neurodegenerative and retinal diseases (57,84–87). These data indicate that the defects caused by pathogenic variants of splicing factors affect multiple retinal cells.

We further identified IFT122 as a differentially spliced transcript with a higher inclusion of the alternative exon 17 in PRPF8-Y2334N organoids. IFT122 is a component of the IFT complex critical for cilia transport and its mutation or loss causes a progressive retinal atrophy in dogs and zebrafish (58–60,88). Alternative inclusion of the same exon 17 has been previously described in a study analyzing RP mutations in another splicing factor, PRPF31 (65). Furthermore, dysfunction of IFT genes has been suggested in the pathogenesis of various forms of RP (62–64). Therefore, cilia represent the primary structural element in RP pathogenesis, with the IFT complex likely playing a central role. The low impact of the Y2334N variant on gene expression observed here aligns with our recent study, in which the same mutation in a homozygous state had a limited effect on RNA expression and splicing in immortalized RPE cells (15). Fluorescence recovery after photobleaching suggested that spliceosomes harboring the Y2334N mutant interact less productively with mRNA than the WT variant (15). This is consistent with our finding that splicing efficiency is reduced for introns with weaker 3′ splice sites, indicating a defect in splice site recognition.

Finally, we observed multiple changes in the expression of circRNAs. It has been previously shown that circRNAs are sensitive to defects in splicing machinery and mutation or downregulation of splicing factors leads to circRNA misexpression (89–93). Similarly, we have observed misexpression of circRNAs in the mouse model expressing the same PRPF8 variant as analyzed in this manuscript (25). Although misexpressed circRNAs differed between the human retina and mouse cerebellum models, our data indicate that alterations in circRNA expression are a common molecular signature of RP mutations in splicing factors. Given their stability compared to other RNA species, misexpressed circRNAs might serve as biomarkers for RP associated with splicing factor mutations. Similarly, misexpression of non-coding RNAs, including circRNAs, has been suggested as a biomarker of retinopathies (94,95). Unfortunately, we could not confirm differential expression of circRNAs in other retinal organoid RP models because the authors sequenced polyA-selected transcripts, excluding circRNA from the RNA-seq analysis (36,65). The unknown function of most circRNAs currently limits our ability to directly correlate changes in individual circRNA levels with the RP phenotype. Nevertheless, their abundance in the human retina (96) indicates a potential functional relevance in this tissue.

In summary, we generated a new retinal organoid RP model for the Y2334N mutation in PRPF8, which reproduces POS defects. The limited changes we detected in the expression and splicing of genes crucial for various retinal cells suggest that a relatively small number of misexpressed genes can drive the RP phenotype. This offers the potential to pinpoint specific genes with defective splicing underlying RP and eventually develop target therapies to correct their splicing.

## Acknowledgement

We would like to thank Jana Machatová-Křížová and Naděžda Vaškovicová for technical assistance. This work was supported by Project JAC CZ.02.01.01/00/22_008/0004575 “RNA for therapy”, co-funded by the European Union, Ministry of Health of the Czech Republic, (NU22-07-00380 to T.B.) and Charles University Grant Agency (1170920 to F.Z.). F.Z, P.B, J.K. and P.K.T are graduate students of Faculty of Science, Charles University in Prague. We also acknowledge the Light Microscopy Core Facility, IMG, supported by MEYS (LM2023050, CZ.02.1.01/0.0/0.0/18_046/0016045 and CZ.02.01.01/00/23_015/0008205), for image acquisition and analysis, and ELIXIR-CZ research infrastructure, funded by MEYS project LM2023055, for bioinformatics analysis.

## Conflict of interest

The authors declare that they have no conflict of interest.

## Supplementary figure legend

**Figure S1.**
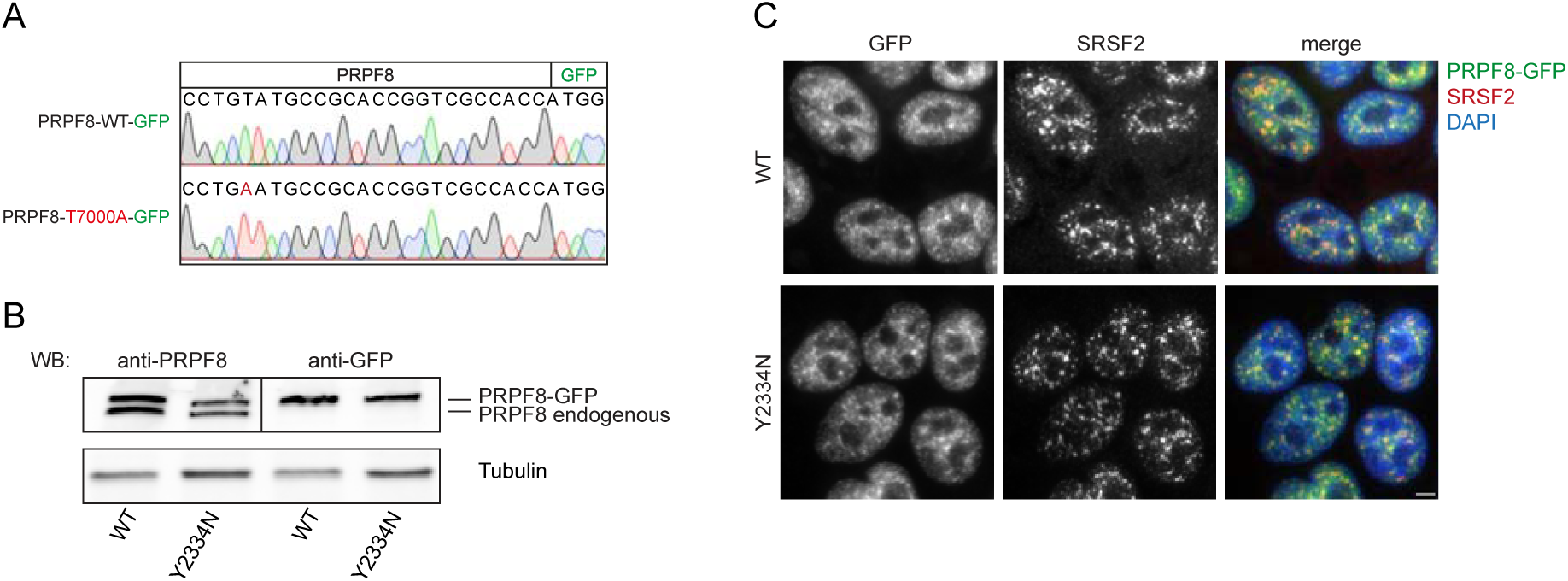
Editing of the PRPF8 gene in human induced pluripotent stem cells (hiPSC). A) Sequence of the GFP-tagged PRPF8 C-terminus of RPPF8-WT and PRPF8-Y2334N hiPSC clones. The c.T7000A substitution in the *PRPF8* gene is highlighted in red. B) Western blot analysis confirming monoallelic editing of the *PRPF8* gene. Reduced endogenous and edited PRPF8 protein levels in Y2334N hiPSC. C) Immunofluorescence analysis reveals that both WT and Y2334N mutant localize to the nucleoplasm and are enriched within splicing speckles as indicated by colocalization with SRSF2. Scale bar = 5 μm.

**Figure S2.**
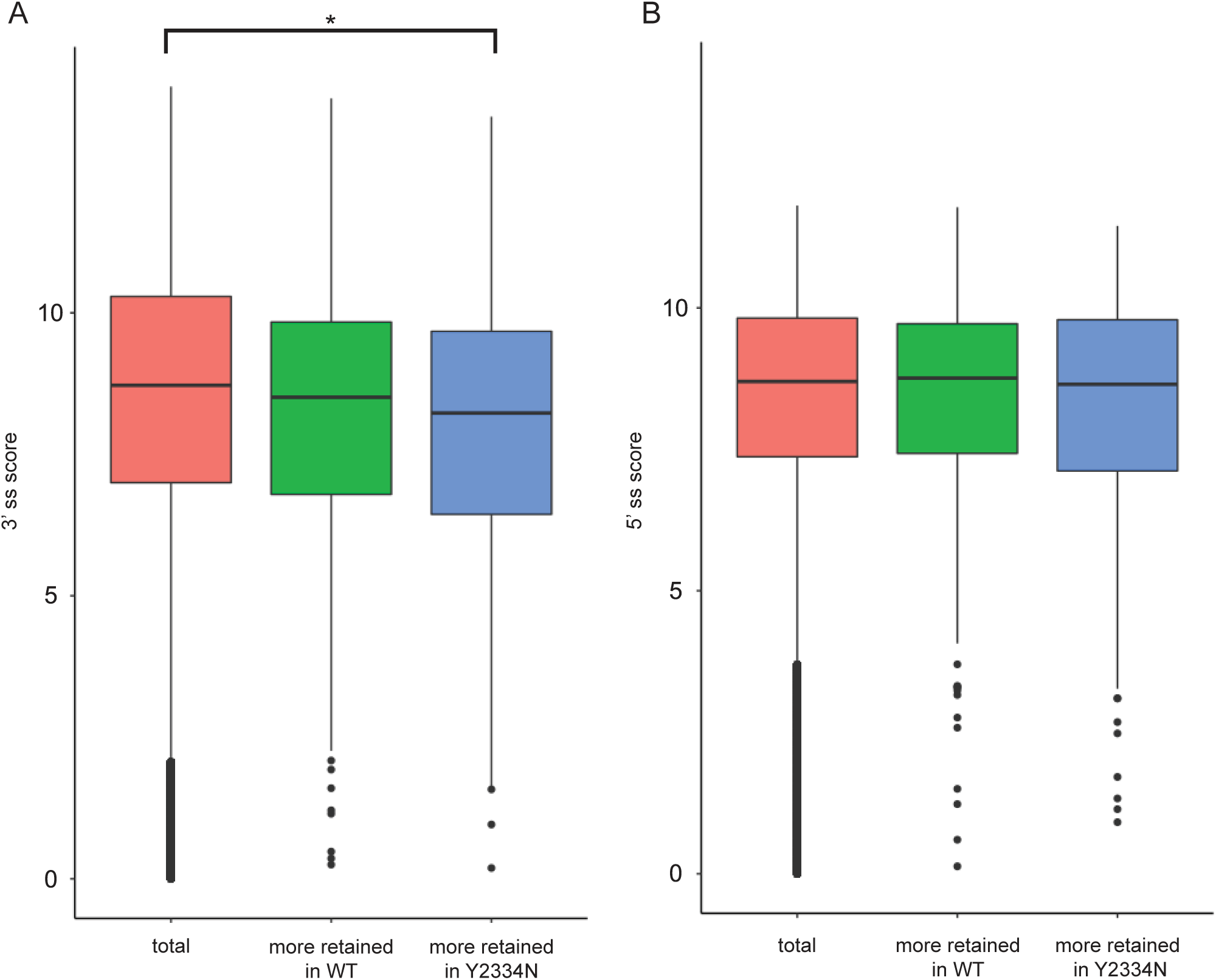
The splice site strength of differentially retained introns is reduced. A) Comparison of the 3’ splice site strength of differentially retained introns in 170-day-old retinal organoids reveals significantly lower strength for introns more retained in Y2334N retinal organoids than the overall human genome average. B) Comparison of the 5’ splice site strength of differentially retained introns reveals no significant difference in the strength of differentially retained introns compared to the overall genome average. Statistical significance was determined using a Wilcoxon rank sum test with continuity correction (* = p < 0.05).

